# Speciation on a complex landscape; phylogeny and demography of Mexican Goodeid fish

**DOI:** 10.1101/2022.12.22.521626

**Authors:** Leeban H. Yusuf, Yolitzi Saldívar Lemus, Peter Thorpe, Constantino Macías Garcia, Michael G. Ritchie

## Abstract

Understanding the phylogeographic history of a group and identifying the factors contributing to speciation is an important challenge in evolutionary biology. Vicariance has clearly played a major role in diversification across diverse groups of organisms, but how this interacts with intrinsic biological features such as changes in reproductive system or sexual selection is less well understood. The *Goodeinae* are a Mexican endemic group of live-bearing fish. Here, we develop genomic resources for species within the *Goodeinae* and use phylogenomic approaches to characterise their evolutionary history. We sequenced, assembled and annotated the genomes of four new *Goodeinae* genomes, including *A. toweri*, the only matrotrophic live-bearing fish without a trophotaenia in the group. We produced a phylogeny of the *Goodeinae* and estimate timings of species divergence. We determined the extent and timing of introgression between the species to assess if this may have occurred during an early radiation, or in more recent episodes of secondary contact. We also analyse patterns in the changes of effective population size for the species and examine the time course of expansion and decline. We used branch-site models to detect genome-wide positive selection across *Goodeinae*, and we specifically ask if this differs in *A. toweri*, where reversal of placental viviparity has recently occurred. We find that Goodeinae diverged rapidly, with reductions in effective population size after each split. Contrary to expectations, we find evidence of gene flow between geographically isolated species, suggesting vicariant speciation was supplemented by limited post-speciation gene flow, potentially as a result of river piracy. Genes under positive selection in the group are likely to be associated with the switch to live-bearing. Overall, our studies suggest that both volcanism driven vicariance and changes in reproductive mode influenced radiation in the Goodeinae.

## Introduction

Understanding the historical biogeography of species groups has traditionally been dominated by vicariance models of speciation (Coyne & Orr 2004; Mayr 1963). Allopatric divergence facilitates the evolution of unique adaptations and other differences, including incompatibilities that can stimulate speciation. In temperate species, this model is thought to involve climatic fluctuations which have patterned both interspecific and more recent intra-specific differences during repeated changes in distribution leading to cycles of allopatry and secondary contact. Many species groups show biogeographic patterns consistent with recent divergence and speciation being patterned by such vicariance, especially during the Pleistocene (Schmitt, 2007; Kadereit and Abbott, 2021), though divergence may have been initiated earlier (Klicka and Zink, 1997; Ebdon *et al*., 2021). Although climatic cycles are perhaps less pronounced outside of the temperate regions, they also occurred in tropical systems that show related patterns of relatively recent allopatric divergence (Thom *et al*., 2020).

Genomic analyses are revealing that many species show recent or historical gene flow, which is compatible with the “repeated allopatry and secondary contact” models of speciation (Harrison and Larson, 2014; Suvorov *et al*., 2022). Some studies suggest that gene flow can facilitate speciation in multiple ways (Marques, Meier and Seehausen, 2019). One of these is when ancient hybridisation can provide genetic material contributing to adaptive radiation in species groups, either by increasing genetic variation or promoting exchange of adaptive genotypes. Some of the most detailed studies of this concern freshwater lacustrine radiations, especially of the African cichlids (Meier *et al*., 2017; Malinsky *et al*., 2018; McGee *et al*., 2020; Svardal *et al*., 2020). Determining the timing and biological significance of gene flow during speciation in species groups with complex evolutionary histories is an important challenge.

Central Mexico is a biodiversity hotspot, but relatively few detailed phylogeographic studies of endemic groups have been completed. Pleistocene climatic fluctuations are often invoked to explain biogeographic patterns, but these can be ‘complex’ (Colin and Eguiarte, 2016). Central Mexico contains much high altitude upland and has been subject to extensive dynamic volcanism during the Miocene and Pleistocene (Ferrari *et al*., 2012). The trans-Mexican volcanic belt is implicated in a Mexican Transition Zone where Nearctic and tropical biotas meet (Morrone, 2010). The highlands may be characterised by sky islands with frequent fragmentation. Mastretta-Yanes *et al*. (2015) argued that such changes may have led to opportunities for hybridisation during speciation of Mexican highland species. Freshwater fish, as well as other organisms (Musher *et al*., 2022) associated with river networks, may have an added complexity to their distributional changes as volcanism has influenced river patterns (Dias *et al*., 2013; Craw *et al*., 2016), including river piracy. In *Poeciliopsis* and *Poecilia genera*,phylogeographic studies implicate a Plio-Pleistocene vicariant event stimulated by the Trans-Mexican Volcanic Belt in driving speciation (Mateos, 2005). Other biogeographical studies show concordant patterns of river vicariance driven by volcanic activity in the Trans-Mexican Volcanic Belt in less well-studied freshwater fish taxa including Goodeins (Domínguez-Domínguez, Doadrio and León, 2006; Domínguez-Domínguez *et al*., 2008; Pérez-Rodríguez *et al*., 2015).

Here we report the first genomic scale studies of the evolutionary history of an endemic group of Mexican fish, the *Goodeinae*. These are a group of viviparous, freshwater fish comprising 36 species across 16 genera. Previous studies have suggested that they radiated in the Miocene ~16 MYA predominantly due to allopatric speciation (Webb *et al*., 2004; Pérez-Rodríguez *et al*., 2015). Their helminth parasites show similar biogeographic patterns (Quiroz-Martínez and Salgado-Maldonado, 2013). It has been proposed that the Goodeins may represent an adaptive radiation following the evolution of live-bearing, and that speciation rate may be influenced by sexual selection. All species within *Goodeinae* are viviparous and matrotrophic, such that offspring depend on a continuous maternal provisioning (Wourms, Grove and Lombardi, 1988; Vega-López *et al*., 2007). Embryos develop extremely rapidly via maternal provisioning through a placental analogue called the trophotaenia (Wourms, Grove and Lombardi, 1988; Hollenberg and Wourms, 1995). Within the *Cyprinodontiformes* (including the *Goodeidae* and *Poecilidae*) speciation rate is higher in live-bearing fish (Helmstetter *et al*., 2016). Interestingly, one species of Goodeid, *Ataenobious toweri*, is thought to have lost the trophotaenia, though little is known about any physiological or genetic changes involved.

Sexual selection has influenced behavioral and morphological sexual dimorphism in the *Goodeinae*. Males cannot inseminate females without female co-operation (Bisazza, 1997), so sexual selection acts strongly on female choice and male secondary sexual traits. Females show preference for male colour patterns across the group (Moyaho, Garcia and Ávila-Luna, 2004; Garcia and Ramirez, 2005; Garcia and Lemus, 2012), particularly in fin structures, and there are pronounced differences in fin size and body shape in males (Garcia, Jimenez and Contreras, 1994; Garcia, Saborío and Berea, 1998; Foster and Piller, 2018). Live-bearing may also give rise to considerable sexual conflict in the group, which can influence speciation rate (Gavrilets, 2000, 2014; Gavrilets and Waxman, 2002). How geography and these intrinsic biological features may interact during speciation is unknown. Previous studies based on microsatellite markers showed that sexually dimorphic species show greater genetic differences between populations (Ritchie *et al*., 2007), but gene flow between populations of a dimorphic species was unrelated to male morphology, instead showing isolation by distance and vicariance (Macías Garcia *et al*., 2012). Mitochondrial DNA sequencing across the group supported evidence for an early radiation, but provided equivocal evidence for a faster speciation rate in more sexually dimorphic species (Ritchie *et al*., 2005).

Here, genomic resources were developed for species within *Goodeinae*, and we used phylogenomic approaches to characterise their evolutionary history. The genomes of four new *Goodeinae* genomes, including *A. toweri* were sequenced, assembled and annotated. We have produced a phylogeny of the species and estimated timings of the divergence. We also analysed the extent and timing of introgression between the species to assess if this may have occurred during an early radiation, or in more recent episodes of secondary contact. We also analysed patterns in the changes of effective population size for the species (many of which are critically endangered) and examine the time course of expansion and decline (Bailey, Garcia and Ritchie, 2007). We used branch-site models to detect genome-wide positive selection across Goodeinae, and we specifically asked if this differs in *A. toweri*, where reversal of placental viviparity has recently occurred.

## Methods

### Read mapping and filtering

Raw genomic reads for each species were trimmed using fastp v.0.20.1 (Chen *et al*., 2018) with default parameters. Cleaned reads were then mapped to a *G. multiradiatus* reference genome (Du *et al*., 2022) using bwa (v. 0.7.17). We used samtools (v. 1.11) (Li *et al*., 2009) to index, sort, mark and remove duplicate reads from each sample and Picard (Institute, 2019) to add read groups to each sample. Subsequently, variant calling was performed on mapped reads using Freebayes (v. 1.3.2) with the following parameters: --report-genotype-likelihood-max --no-population-priors --use-best-n-alleles 4 --hwe-priors-off --use-mapping-quality --ploidy 2 --theta 0.02 --haplotypelength −1 --genotype-qualities. Finally, variants were filtered using GATK hard-filtering best practices guidelines (Van der Auwera *et al*., 2013) (QD < 2.0, QUAL < 30.0, SOR > 3.0, FS > 60.0, MQ < 40.0, MQRankSum < −12.5, ReadPosRankSum < −8.0).

### Phylogenetic history and dating of Goodeinae

To infer the phylogenetic history of the *Goodeinae*, we further filtered variants used in our phylogenetic analysis. We removed variants with missing genotypes across any of our samples, non-biallelic positions, and positions with either no alternative allele or only alternative alleles using bcftools. Additionally, we sample SNPs equally across the genome and pruned these by sampling 1 SNP for every 100 SNPs, to try to select independent histories. We used the following parameters in bcftools: bcftools +prune -w 100bp –n 1. The pruned and filtered VCF was then converted into a phylip alignment by concatenating all variants using vcf2phylip (Ortiz, 2019) for all nine species. We first inferred phylogenetic relationships using a IQTREE2 (v. 2.1.4) using ModelFinder and 1000 bootstraps. Since maximum-likelihood concatenation approaches may fail when incomplete lineage sorting is prominent, we sought to infer a species tree using SVDquartets (Chifman and Kubatko, 2014, 2015) in paup (v.4.0a)(Wilgenbusch and Swofford, 2003) with 100 bootstraps and using filtered, pruned variants.

To infer divergence times and effective population sizes for ancestral nodes, we used the A00 model in BPP (v4.4.0)(Flouri *et al*., 2018) and putatively selectively-neutral non-coding loci. To generate our selectively-neutral non-coding dataset, we first estimated substitution rates of four-fold degenerate sites (non-conserved regions) in 21 species across *Cyprinodontiformes* to identify regions with conserved substitution rates using PhastCons (Siepel et al. 2005; Pollard et al. 2010; Yusuf et al, in prep). Variants falling within these conserved regions were subsequently removed from our dataset, alongside coding regions, using bedtools (v. 1.12) (Quinlan and Hall, 2010). Additionally, to ensure sampled loci represented different evolutionary histories as required by BPP, we made 10kb windows that were at least 100kb away from each other. This yielded 2,740 loci of variable size that were converted into alignments with no missing sites for any species using vcf2phylip (Ortiz, 2019). To estimate divergence times and effective population sizes, we used A00 analysis with default settings except for amendments to theta (0.003) and tau (0.03) priors. We used a burn-in period of 8,000 and 100,000 samples. Divergence time estimates and effective population size estimates were scaled to geological time using the *Xiphophorus maculatus* mutation rate (3.5 × 10^-9^) from Schumer et al. (2018) and a generation time of 1 via R package bppr (v.0.6.1) (Angelis and Dos Reis, 2015).

### Principal component analysis

To supplement the phylogenetic analysis, we sought to understand sample relationships using principal components analysis. We utilized hard-filtered biallelic SNPs that contained no missing data across our samples and were pruned based on distance to nearest SNP. We used plink2 to compute principle components analysis from 8,161,340 SNPs using the following parameters: “--double-id --allow-extra-chr -- set-missing-var-ids @:# --vcf-half-call m –pca” (Frazer *et al*., 2007; Chang *et al*., 2015).

### Demographic inference and population size changes

In order to detect recent changes in effective population size for each species from a single representative sample, we used the pairwise sequentially Markovian coalescent (PSMC) model (Li and Durbin, 2011). We converted mapped reads for each species individually into mpileup format using samtools and bcftools with the -c option (Li *et al*., 2009). We then filtered out variants with depth lower than 10, a depth higher than 100 and mapping quality lower than 30, and converted variants into *fastq* format using the *vcf2fq*function of the vcfutils.pl program available via samtools (Li *et al*., 2009). To determine choice of time interval parameters, we tested a range of interval parameters and found no difference in our results. As a result, we used default interval parameters of 4+25*2+4+6 for our final analysis. To scale estimates of effective population size, we used the *Xiphophorus* maculatus mutation rate (3.5 × 10^-9^) from Schumer et al. (2018) and a generation time of 1.

To understand how changes in population size affected genetic divergence genomewide, we computed absolute nucleotide divergence (D_XY_) in 50 kb non-overlapping windows between three relatively closely-related pairs: *A. splendens* and *X. captivus*; *I. furcidens* and *X. resolanae;* and *G. multiradiatus* and *A. toweri*, using the program pixy (v. 1.2.3.beta1) (Korunes and Samuk, 2021).

### Testing global and local patterns of introgression

To test for evidence of introgression between species, we calculated Patterson’s D-statistic using the *Dtrios* function in Dsuite v.4 (Green *et al*., 2010; Malinsky, Matschiner and Svardal, 2020) with hard filtered variants (described above). Specifically, we inferred introgression across 56 trios with trios arranged according to phylogenetic relationships in our inferred species tree. As well as calculating D-statistics for all 56 trios, we also calculated the admixture fraction (*f4-ratio*). To control for false discovery, we applied a Benjamini-Hochberg procedure to p-values for all trios and kept only trios with corrected p-values lower than 0.05 (Benjamini and Hochberg, 1995). To filter out trios with unreliable signals, we removed trios where *D* and the *f4-ratio* was lower than 1%.

Because introgression signals can represent both (a) very recent introgression between extant lineages and (b) ancient introgression between ancestral lineages, we inferred the *f-branch* statistic via Dsuite. The *f-branch* statistic computes allele sharing using *f4-ratios* between P3, where P3 is an outgroup species to P1 species and P2 species, and the descendants of a branch labelled *b* (i.e., the descendants of P2, for example) relative to a sister branch labelled *a*. This statistic allows for correlated *f4-ratios* to be disentangled and specific branches with strong signals of recent introgression to be identified.

To supplement tests for introgression, we also sought to understand local patterns of introgression and identify genes that may have been introgressed. We estimated fd genome-wide in 150bp windows using the *Dinvestigate* program in *Dsuite* (Martin, Davey and Jiggins, 2015; Malinsky, Matschiner and Svardal, 2020). Specifically, we surveyed introgression genome-wide only for trios that showed the most consistent signals of introgression across all genome-wide estimates of introgression. To determine genes which may have introgressed across trios, we subsetted our dataset to only include windows within the top 10% of windows based on f_d_ genome-wide. We then used the *GenomicRanges* package (Lawrence *et al*., 2013) in R to determine whether any windows overlapped with annotated genes in our gff file. This subsetting approach allowed for the conservative identification of high-confidence introgressed genes maintained across multiple independent introgression events or an ancient event(s).

### Detecting positive selection in rapid diversification of *Goodeinae*

To test for positive selection in short internal branches leading to the rapid diversification of Goodeids, we used the reference genome from *G.multiradiatus* and our hard-filtered variant calls to produce a consensus sequence alignment for each gene using the vcf2fasta software (https://github.com/santiagosnchez/vcf2fasta). To ensure gene alignments did not contain de-novo genes not found in our outgroup species *C.baileyi*, alongside the reference genome and hard-filtered VCF file, we provided a filtered gene annotation file containing only annotated genes found in both *G. multiradiatus* and *C. baileyi*. To produce this consensus gene annotation, we lifted annotations from *C. baileyi* to the *G. multiradiatus* annotation; out of 29,739 genes we considered only 20,138. Nucleotide gene alignments were then aligned using MACSE v2.05 and the following parameters: “-prog refineAlignment -optim 2 -local_realign_init 0.001 -local_realign_dec 0.001”. Nucleotide alignments were then masked for any remaining internal stop codons and frameshift mutations using the parameters: “-codonForInternalStop NNN -codonForInternalFS --- -charForRemainingFS – ”. Finally, nucleotide alignments were translated into amino acid alignments and were subsequently used to produce a codon alignment using the program PAL2NAL and the parameters: “-nogap -nomismatch”.

We used two approaches to detect positive selection. First, for each internal branch and codon alignment, we asked whether a proportion of sites were evolving under positive selection relative to background branches using aBSREL (Smith *et al*., 2015). For each codon alignment, we used the inferred species tree phylogeny. Since all internal branches were tested separately, p-values were corrected for multiple testing and filtered (p < 0.05) using the Benjamini-Holm procedure. Secondly, for all internal branches considered together, we tested whether specific codons in each codon alignment (gene) were under positive selection in internal branches, using the codon model MEME (Murrell *et al*., 2012) and the species tree phylogeny. Again, p-values were corrected for multiple testing using the Benjamini-Hochberg procedure (Benjamini and Hochberg, 1995). Extraction of results for aBSREL and MEME from JSON files was performed using Python package *phyphy* ( J. Spielman 2018). Additionally, we performed gene ontology via R package ‘clusterProfiler’ on genes found to have evidence of positive selection on at least one internal branch (Yu *et al*., 2012).

To facilitate understanding of how the trophotaenium was lost in *A. toweri*, we performed a branch-site test via aBSREL on the terminal branch leading to *A. toweri*. We supplemented this test for positive selection in *A. toweri* in two ways. First, by overlapping positively selected genes with genes upregulated in the trophotaenia of *G.multiradiatus*, taken from a recent genomic and transcriptomic analysis of *G.multiradiatus* (Du *et al*., 2022). Second, we used the Ensembl Variant Effect Predictor (VEP) tool to infer, using genome annotation information, high-impact mutations in protein-coding genes that may drastically alter protein function (McLaren *et al*., 2016). Altogether, we searched for genes upregulated specifically in the trophotaenium of *G. multiradiatus* that may also have high-impact mutations in *A. toweri* indicating potential pseudogenization.

## Results

### Rapid diversification of Goodeinae occurred in the middle Miocene

Both approaches to inferring the phylogeny gave a consistent species tree topology with complete bootstrap support and complete posterior probabilities, and recapitulate relationships inferred in previous molecular phylogenies produced using mitochondrial sequences (Webb *et al*., 2004). We find that following the divergence of *Empetrichthyinae* (represented by *C. baileyi*), a group containing *A. toweri* and *G. multiradiatus* diverged next, supporting previous suggestions that the evolutionary reversal of trophotaenia development occurred in *A. toweri*, as opposed to an ancestral placement of *A. toweri* at the crown of the *Goodeinae*. We resolve the placement of *C. lateralis*, which diverged following the split of *A. toweri* and *G. multiradiatus*. The remaining species fall into two groups; a group containing *I. furcidens* and *X. resolanae*, and another containing *G. atripinnis*, *A. splendens* and *X. captivus*, both of which are concordant with previous phylogenetic reconstructions of *Goodeinae* (Webb *et al*., 2004) (Figure 1). These patterns are also recovered in principal components analysis of filtered SNPs (Supplementary figure 1). Whilst PC1 likely reflects technical variation separating *G. multiradiatus* from all other species, PC2 and PC3 cluster species reflect relationships described in the phylogenetic inference.

**Figure 1:**
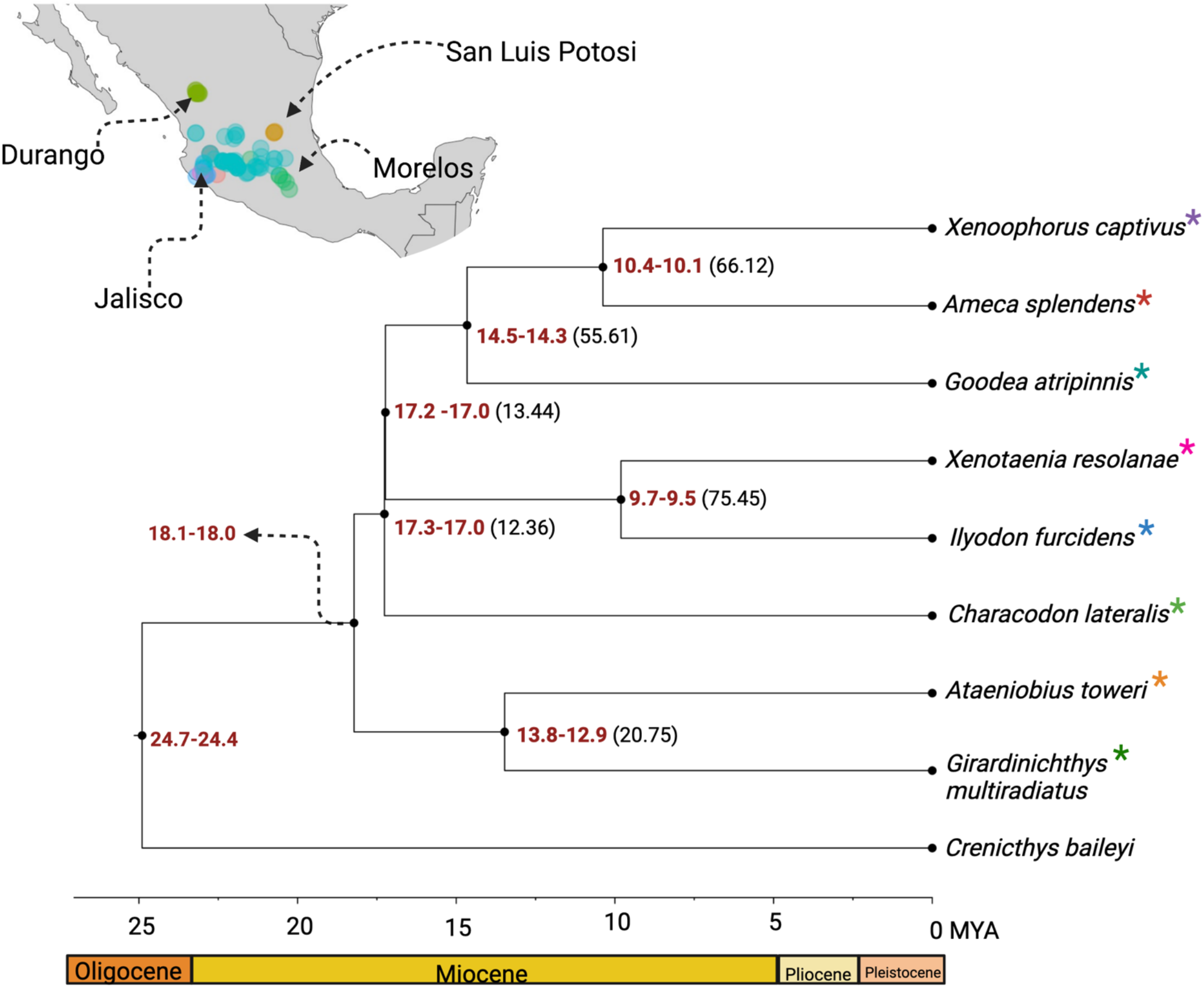
Species tree of the family *Goodeidae*. Scaled divergence times are shown for each node (in red) and gene concordance factors are shown for relevant nodes (in brackets). Colours in asterisks on the phylogeny denote species in map showing distribution of species. Distribution information taken from FishNet2.net.

Since classic measures of phylogenetic support tend to overestimate confidence in phylogenomic datasets (Kumar *et al*., 2012; Salichos and Rokas, 2013; Yang and Zhu, 2018), we inferred gene concordance factors using the species tree inferred with ASTRAL and 19,908 gene trees via IQTREE2. Gene concordance factors quantify the percentage of gene trees that contain a given branch in the species tree. We find that almost all branches show extremely low gene concordance, except for branches leading to the split of *G. atripinnis*, *A. splendens* and *X. captivus*, and of *I. furcidens* and *X. resolanae* (Figure 1). This gene tree discordance is confirmed by inference of normalized quartet scores in ASTRAL where only 60% of quartets across all gene trees are also found in the species tree. Additionally, around the crown radiation, we observe short branch lengths and broad variation in support for alternate topologies, indicating frequent incomplete lineage sorting, presumably as a result of rapid speciation of *Goodeinae* (Figure 1).

To understand when diversification of *Goodeinae* began, we estimated divergence times and effective population sizes using filtered dataset of 2,740 putatively neutral non-coding loci, the species tree estimated above using ASTRAL and a multi-species coalescent approach. We find that crown age of the group and the divergence of *Empetrichthyinae* occurred between 24.7 – 24.4 MYA. We estimate that the diversification of *Goodeinae* began 18 MYA with the diversification of a clade containing *G. multiradiatus* and *A. toweri*. Following the split of *G. multiradiatus* and *A. toweri*, divergence of two clades containing (a) *G. atripinnis*, *A. splendens* and *X. captivus*, and (b) of *I. furcidens* and *X. resolanae* occurred alongside divergence of *C.lateralis* around 17.4 – 17.2 MYA. We estimate that the split of *C.lateralis* from the others was accompanied by considerable reduction (~100 fold) in effective population size, likely owing to vicariance as a result of hypothesized increase in volcanism (Webb *et al*., 2004). This is supported by similar reduction (~5 fold) in effective population size following the split of *C. baileyi*. Divergence of *A. toweri*, where evolutionary reversal of trophotaenia is hypothesized, occurred around 13.9 – 13.1 MYA. Altogether, our divergence time and effective population size estimates suggest incomplete lineage sorting is likely explained by rapid vicariant speciation events that occurred early in the diversification of Goodeids.

### Limited evidence of introgression supports vicariant speciation of *Goodeids*

Alongside incomplete lineage sorting, gene flow may also produce patterns of underlying gene tree conflict with the species tree. To assess the potential importance of gene flow in explaining patterns of genetic variation in Goodeids, we computed Patterson’s D statistic and the f4-ratios for all possible species combinations, without assuming or arranging species according to their species tree relationships. This approach produces conservative estimates of allele sharing (D_min_) for all trios across all possible topologies (Malinsky *et al*., 2018; Malinsky, Matschiner and Svardal, 2020). We find that 41 out of 56 tested trios (73%) have a significantly positive D-statistic value (D_min_) after correcting for multiple testing, with average genome-wide allele sharing of 4.7% (Figure 2).

**Figure 2:**
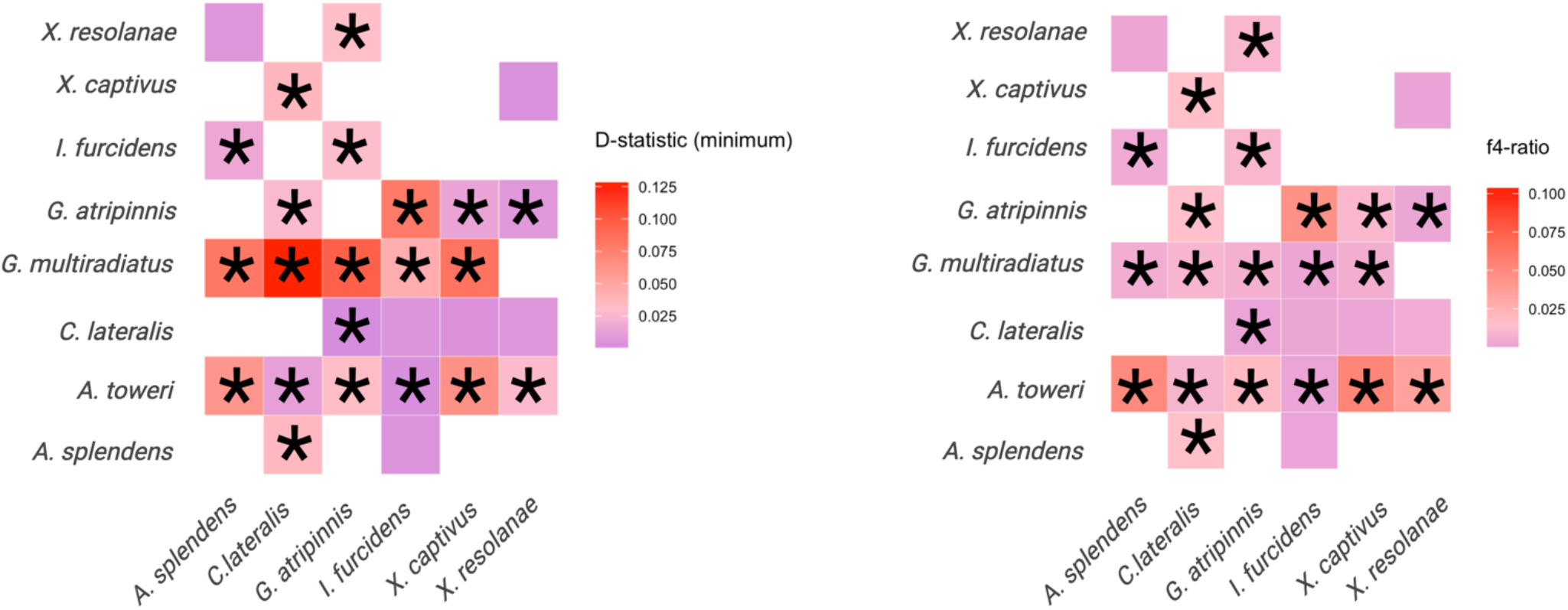
Evidence of gene flow between species in *Goodeinae*. Plots showing D-statistics and f4-ratios in a pairwise manner. Significant allele sharing after multiple-testing correction is indicated by panels with an asterisk.

However, estimates of introgression are not independent since a single, ancestral gene flow event may result in erroneous interpretations of recent, widespread gene flow. We therefore re-calculated Dmin by reorganising trios according to the relationships observed in the species tree inferred using ASTRAL. Specifically, we calculated the f-branch statistic *f*b(C) which utilises correlated f4-ratios to provide branch-specific estimates of introgression. We find support for gene flow events (*f*b(C) > 10%) between *A. toweri*, which is endemic to San Luis Potosí, and all other species apart from *G. multiradiatus*. Additionally, we find support for weaker introgression between *G. multiradiatus* and an ancestral branch leading to the divergence of *G. atripinnis, X. captivus* and *A. splendens*. Altogether, we suggest there is evidence for a potential ancient gene flow event between the ancestor of *G. multiradiatus* and *A. toweri* and an ancestor of *G. atripinnis, X. captivus* and *A. splendens*.

To examine patterns of local introgression and identify introgressed genes, we computed fd for the introgression event(s) with most support. We examined introgression between trios containing *G. multiradiatus* (P1), *A. toweri* (P2) and all other Goodeines (P3), with *C. baileyi* (P4) used as the outgroup (Supplementary Figure 2). We find similar levels of mean admixture proportions across surveyed across comparisons, with highest levels of mean admixture (~7.4%) observed in *G. atripinnis*, which shares its range with *A. toweri*, and in *X. resolanae*, which is currently allopatric from *A. toweri*. In *C. lateralis, A. splendens* and *X. captivus*, lower levels of genome-wide admixture with *A. toweri* are observed (5.7 – 5.9%), perhaps owing to differential loss of introgressed variation.

Across all comparisons, we find 64 genes that show evidence of high f_d_ (top 10%) across all trios. Among those, we find evidence of introgression in class II, major histocompatibility complex, transactivator (CIITA), which mediates expression of other MHC II genes and is under divergent sequence evolution in other viviparous lineages (Roth *et al*., 2020). We found a significant enrichment of genes related to sugar and fat metabolic processes, as well as activation and regulation of transcription factors (Supplementary table 1). However, gene ontology analysis of introgressed genes maintained across all comparisons showed no significant enrichment of any biological processes, after correcting for multiple testing. Since introgressed genetic variation may be differentially lost by drift or removed by purifying selection, we also conducted gene ontology analysis on the comparison (introgression between *A. toweri* and *X. resolanae*) with most introgressed genetic variation to recover as much genetic variation that was initially exchanged as possible. For this comparison, we found significant enrichment of genes involved in cilium projection and organization, cell projection and gastrulation (Supplementary table 2).

### Evidence for large changes in population size across Goodeid diversification and on contemporary timescales

To identify contemporary changes in contemporary effective population size for each species following their divergence, we reconstructed the demographic trajectory through time for 9 species in *Goodeinae* including *C. baileyi* using the Pairwise Sequentially Markovian Coalescent (PSMC) approach (Li and Durbin, 2011). We find that most species showed a coincident large increase in population size starting around 5 MYA (well after their initial radiation), except for *G. multiradiatus* and *X. captivus*, which began diverging around 1 MYA. Populations then declined progressively since the middle Pleistocene, though there is substantial variation in the initiation and rate of decline (Figure 3). Decline was persistent until recently, with only *G. atripinnis* and to a lesser extent *G. multiradiatus* showing subsequent growth then decline in more recent times, prior to the Holocene. Altogether, our analyses indicate considerable, recent fluctuation in effective population size in *Goodeinae*, paired with similar fluctuations in effective population size during early diversification of the family.

**Figure 3:**
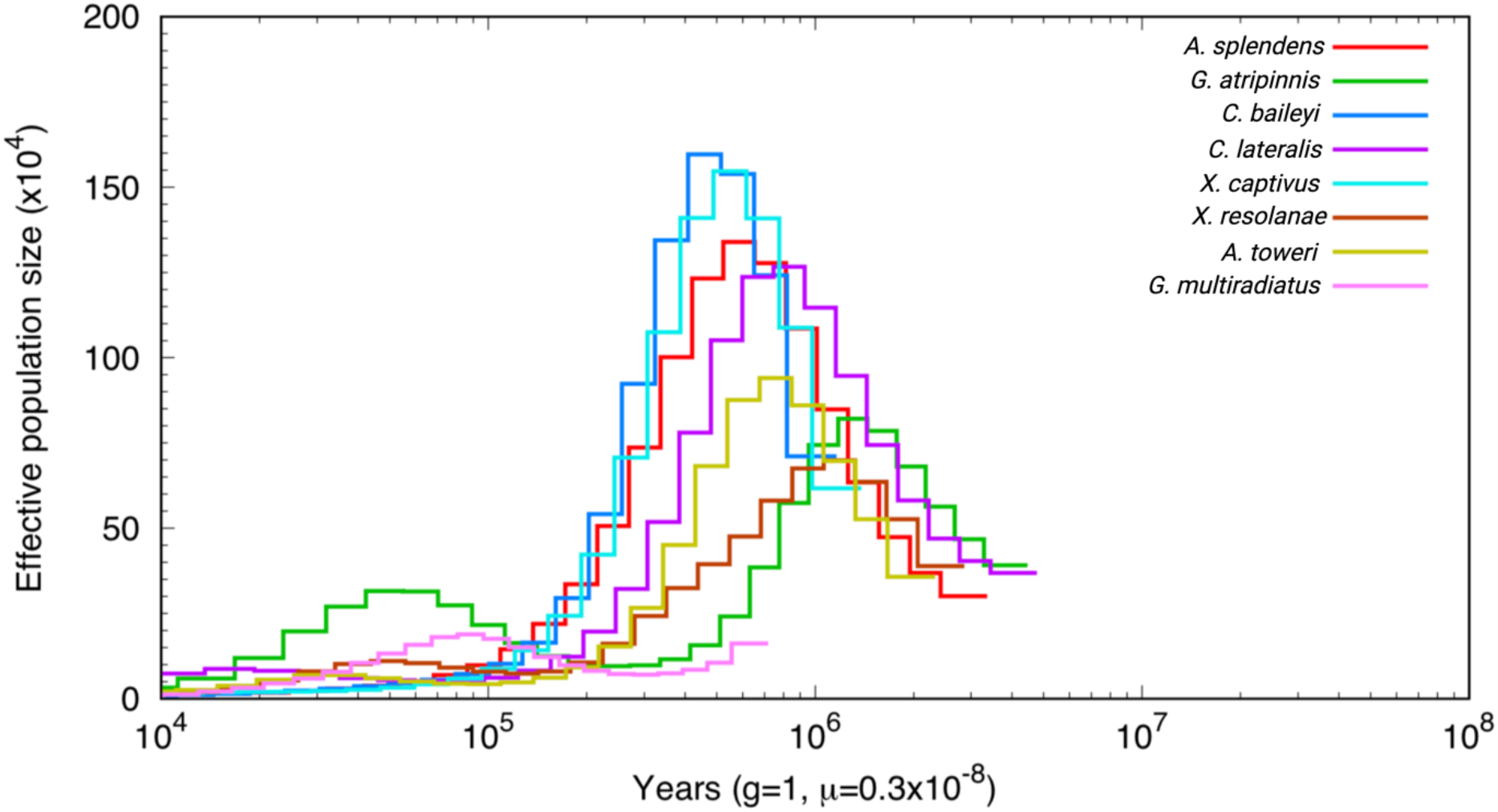
Concordant changes in contemporary effective population size across *Goodeidae*. Inferred effective population sizes through time using pairwise sequentially Markovian coalescent (PSMC) analysis. The X axis indicates scaled time estimates, assuming a mutation rate of 3.5 × 10^-9^ and a generation time of 1.

To assess the potential impact of changes in effective population size on genome divergence between closely related species, we calculated absolute nucleotide divergence (D_XY_) in autosomes and the X chromosome between closely related species in Goodeids. An important consequence of population contraction is that genetic variation is differentially affected on the X chromosome compared to the autosomes. Specifically, ancestral population contractions may lead to reduction of genetic diversity on the X chromosome compared to autosomes, resulting in lower levels of D_XY_ on the X chromosome (Van Belleghem *et al*., 2018). However, since D_XY_ measures nucleotide differences accrued since the divergence time, it is not affected by contemporary changes in population size after species divergence. We find significantly lower D_XY_ on the X chromosome compared to the autosomes between *I. furcidens* and *X. resolanae* (*Welch’s t* = 3.1389, df = 92.575, p-value < 0.001), and between *G. multiradiatus* and *A. toweri* (*Welch’s t* = 4.7094, df = 87.64, p-value < 0.001), but not between *A. splendens* and *X. captivus* (*Welch’s t* = 1.7081, df = 92.675, p-value = 0.09), though we did also observe lower levels of divergence on the X chromosome in this comparison also (Figure 4).

**Figure 4:**
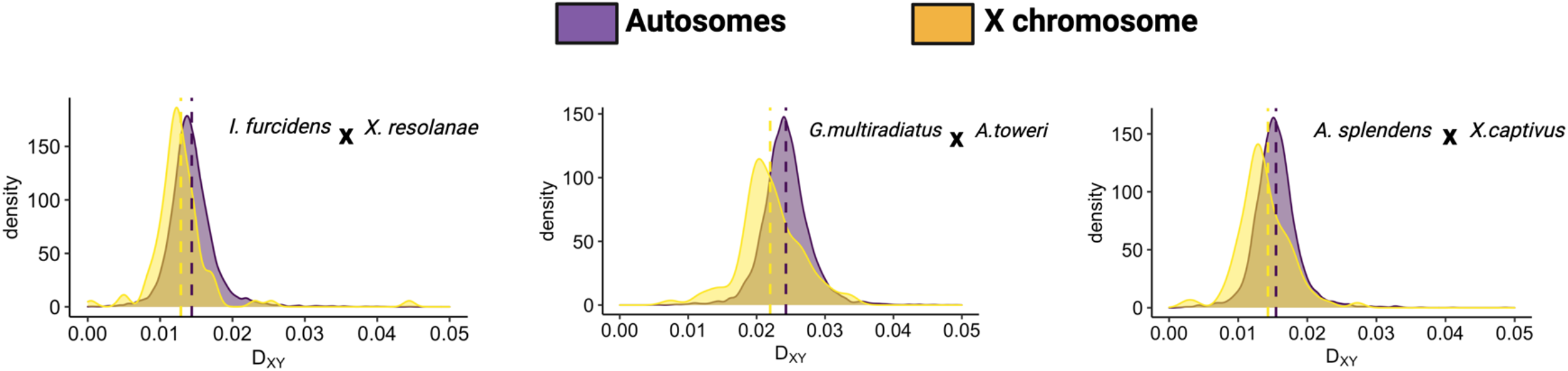
Differences in absolute divergence between autosomes and the X chromosome in three comparisons of closely related species in *Goodeinae*. Distributions of absolute nucleotide divergence (D_XY_) calculated in non-overlapping 50kb windows. Coloured dotted lines show means for both autosomes and X chromosome for each pair.

### Positive selection across *Goodeinae*

To understand what genes may have been associated with diversification across *Goodeinae*, we performed branch-site tests in all internal branches in the species tree. After multiple testing correction, we found 1482 genes (7.3% of all tested) that were under positive selection in at least one internal branch. Of those genes, 1293 (6.4%) of these are only under positive selection in a single branch, with 168 (0.8%) showing evidence of positive selection in two branches and 21 (0.1%) genes evolving under positive selection in three branches. Genes under selection across three internal branches include canonical immune response genes like interleukin-2 receptor subunit gamma (*IL-2RG*), cluster of differentiation 4 (*CD4*) and Lymphocyte activation gene 3 (*LAG3)*. Gene ontology across all positively selected genes (1482) shows a significant enrichment (FDR < 0.05) of genes involved in DNA damage and repair, reproduction and cilium movement, among others (Figure 5; Supplementary table 3).

**Figure 5:**
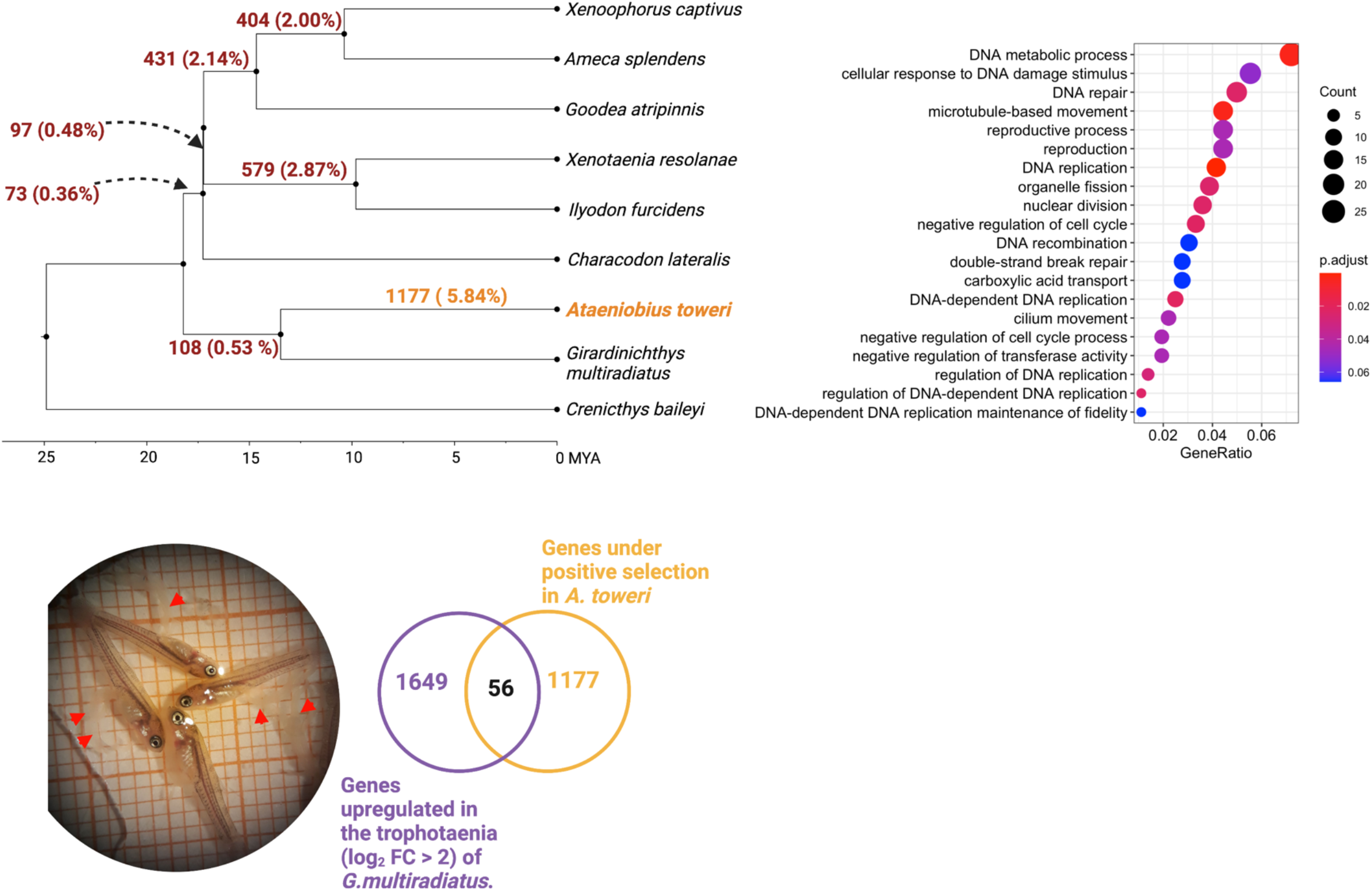
Testing for evidence of positive selection in branches across *Goodeinae*. A) Phylogeny of *Goodeinae* showing the number of genes and the percentage of genes under positive selection at tested branches. B) Gene ontology terms associated with genes under positive selection across *Goodeinae*. Colours denote adjusted p-value (for multiple testing) and size of data point denotes number of under positive selection. GeneRatio indicates the ratio of the number of genes under positive selection and the background gene set for each enrichment term. C) Pregnant *G. multiradiatus*. Red lines point towards exposed trophotaenia. Venn diagram shows the number of genes upregulated in the trophotaenia of *G. multiradiatus*, the number of genes under positive selection in *A. toweri*, and their overlap.

Additionally, we examined the number and identity of genes under positive selection in each internal branch. In the branch leading to the split of *G. multiradiatus* and *A. toweri*, we find 108 genes under positive selection. In subsequent internal branches leading to the split of *C. lateralis* and the ancestor of *I. furcidens, X. resolanae, G. atripinnis, A. splendens* and *X. captivus*, we find even fewer genes under positive selection (73 and 97, respectively). We detect many more genes under positive selection in longer internal branches leading to the split of *G. atripinnis* from *A. splendens* and *X. captivus* (431 genes), *A. splendens* and *X. captivus* (404 genes), and *I. furcidens* and *X. resolanae* (579 genes). These include genes presumed to be important to the evolution of viviparity such as follicle stimulating hormone receptor (*FSHR*), androgen receptor (*AR*), vascular endothelial growth factor a (*VEGFA*), insulin growth factor 1 receptor (*IGF1R*) and insulin-degrading enzyme (*IDE*).

### Identifying genes associated with evolutionary reversal of trophotaenia development in *Ataeniobius toweri*

Based on the phylogenetic inference above and morphological evidence, the trophotaenia was likely lost in *A. toweri* following its split from *G. multiradiatus*. We therefore performed additional branch-site tests for positive selection on the terminal branch leading to *A. toweri*. Altogether, 1177 genes were found to be under positive selection here, after correcting for multiple testing. We found no enrichment of any gene ontology terms for genes under positive selection in *A. toweri*. We overlapped these with genes upregulated (log_2_ fold change > 2) in the trophotaenia compared to all other tissues of *G. multiradiatus*. We found 56 genes which showed significant upregulation in trophotaenia and under positive selection in *A. toweri*. Whilst gene ontology analysis showed no significant enrichment of any terms after multiple testing correction, we note that non-significant terms relate to fin and appendage development (adjusted p-value = 0.13), perhaps consistent with a potential role for these genes in trophotaenia developmental reversal (Supplementary table 4).

We also used Variant Effect Predictor (VEP) analysis to detect high-impact mutations that may have disruptive effects on protein structure or function in *A. toweri*. Specifically, we searched for homozygous high-impact mutations specific to *A. toweri* and overlapped these with the 56 trophotaenia-specific genes under positive selection in *A. toweri*. Only 6 genes showed evidence high impact mutations, including a *cadherin related family member 5* (CDHR5), which modulates assembly of the intestinal brush border (Crawley *et al*., 2014), and *Fraser extracellular matrix complex subunit 1* (FRAS1), which is involved in with epithelial-mesenchymal formation during embryonic development (Pavlakis, Chiotaki and Chalepakis, 2011).

## Discussion

Understanding the spatial context of speciation and identifying the factors contributing to this is an important challenge in evolutionary biology. Since demography and evolutionary history interact with evolutionary forces, inferring species relationships alongside estimates of demography is helping to resolve the relative importance of different evolutionary processes. In *Goodeinae*, the development of live-bearing leads to females provision offspring throughout development and males contribute fewer resources, setting the stage for sexual conflict, and there are varying levels of sexual dimorphism observed throughout the clade. Live-bearing has influenced speciation rates in the Cyprinodontiformes (Helmstetter *et al*., 2016). Analyses of sexual dimorphism and speciation rate in the Goodeinae have provided ambiguous results (Ritchie *et al*., 2005, 2007). Extrinsic factors, primarily recurrent volcanic activity throughout Central Mexico, have also been proposed as an important driver of speciation rates in *Goodeinae*. Our results provide detailed insight into the evolutionary history of *Goodeinae* and suggest a joint role for vicariant speciation with gene flow, and the importance of matrotrophic viviparity in the diversification of *Goodeinae*.

Overall, we found support for rapid divergence early on in the evolutionary history of *Goodeinae*, largely confirming previous phylogenetic analyses (Webb *et al*., 2004; Ritchie *et al*., 2005; Dominguez-Dominguez *et al*., 2010). We find that diversification of *Goodeinae* began 18 MYA with the split of *Ataeniobius* and *Girardinichthys* and was followed by rapid divergence of *Characodon* and lineages including *Goodea*, *Xenotaenia, Xenoophorus, Ilyodon* and *Ameca* around 17 MYA. One noticeable change in our tree is that *C. lateralis* is no longer basal, as seen in trees based on mtDNA (Webb *et al*., 2004; Dominguez-Dominguez *et al*., 2010) but within the main body of the *Goodeinae*. The position of *C. lateralis* is supported in both of our independent coalescent analyses (SVDquartets and ASTRAL) of whole-genome sequence data, despite underlying gene conflict. However, biogeographic analyses have supported an earlier branching off of the Characodontini (Dominguez-Dominguez *et al*., 2010), so this may need to be reconsidered in future.

These radiations coincide with the beginning of activity in the Trans-Mexican Volcanic Belt, and are consistent with previous phylogenetic analysis and fossil evidence (Webb *et al*., 2004; Ferrari *et al*., 2012). Our results disagree with previous analysis supporting basal divergence of *Characodon* in Durango, and instead support diversification of *Goodeinae* that likely occurred east to west with divergence of *Ataeniobius* and *Girardinichthys* (Webb *et al*., 2004). Whilst we find complete statistical support for these early divergence events, we also find extensive underlying gene tree conflict, with only 12-13% of protein-coding gene trees supporting these branches and most gene trees supporting polyphyletic relationships. Here, gene tree conflict is expected to be generated by incomplete lineage sorting as a result of the rapid early diversification. Divergence of *Goodea, Xenotaenia, Xenoophorus, Ilyodon* and *Ameca* occurred 10-9 MYA, likely caused by another bout of volcanic activity beginning in the late Miocene (11 MYA)(Ferrari *et al*., 2012). Altogether, the phylogeographic history of *Goodeinae* inferred here suggests early divergence of the group was characterised by repeated volcanic events occurring throughout the Miocene facilitating bursts of vicariant speciation, as opposed to potential idiosyncratic, intrinsic factors specific to each divergence event (Webb *et al*., 2004; Ritchie *et al*., 2005; Crisp and Cook, 2007).

Whilst geological changes likely contributed to early diversification of *Goodeinae*,recurrent volcanic activity may have also facilitated secondary contact and gene flow. Recent work has shown that classic models of vicariant speciation may be relatively simplistic and obscure more complex speciation histories (Kopuchian *et al*., 2020; Naka and Pil, 2020). Here, we examined evidence of introgression between species in *Goodeinae*, characterizing the degree of allele sharing genome-wide and at particular genomic regions. We found limited evidence of introgression between *A. toweri* and species within *Goodea*, *Xenotaenia*, *Xenoophorus*, *Ilyodon* and *Ameca*, which likely represent a single gene flow event between an ancestral lineage leading to the diversification of the aforementioned species and the ancestor of *A. toweri* and *G. multiradiatus*. This is partially supported by biogeographical evidence from multiple species of *Goodeinae* likely co-occurring with *A. toweri* in the tributaries of Rio Panuco at some point in their evolutionary history (Webb *et al*., 2004). Some previous analyses of genetic differentiation between populations of dimorphic and monomorphic *Goodeinae* species found gene flow to be occurring, but less readily between populations of monomorphic species (Ritchie *et al*., 2007). Recent genetic analysis of populations within *Goodea* and *Ilyodon* suggest connectivity and gene flow between species and populations is highly variable and mediated by complex topographical changes (Beltrán-López *et al*., 2017, 2021). Whilst we detect moderate levels of ancient gene flow, it is not possible here to tell whether gene flow is adaptive and has contributed to phenotypic diversity and diversification, as observed in other systems (Malinsky *et al*., 2015; Meier *et al*., 2017; Svardal *et al*., 2020), but some of these introgressed genes were maintained.

Changes in contemporary and ancient population size throughout the evolutionary history of *Goodeinae* may be common, and several species of the group are endangered. We found dramatic concordant changes in effective population size (N_e_) for almost all species in *Goodeinae* around 500 KYA, followed by concordant and steady reduction in effective population size in all species. This is largely consistent with distributions and the current conservation status of Goodeinae found here (Bailey, Garcia and Ritchie, 2007; Lyons *et al*., 2019). However, it is important to note that absolute time estimates should be interpreted with caution given mutation rates from *Xiphophorus* that were used to scale parameter estimates in this study may be different, and mutation variation within Goodeids may vary. Alongside changes in contemporary changes in effective population sizes, we also find considerable reduction in population size following the split of *C. baileyi* and *C. lateralis*, suggesting possible vicariance as a result of volcanic activity resulted in uneven population sizes in ancestral lineages. These results are further supported by greater genetic divergence observed in autosomes compared to X chromosomes in three comparisons of closely related Goodeinae species. Ordinarily, genetic divergence is expected to be greater on the X chromosome than on the autosomes due to a disproportionate role of the X chromosome in driving reproductive isolation (Meisel and Connallon, 2013; Charlesworth, Campos and Jackson, 2018). However, marked differences in levels of genetic divergence between the X chromosome and autosomes may also be explained by differences in effective population size (Meisel and Connallon, 2013; Wolf and Ellegren, 2017). For example, when absolute genetic divergence between closely-related species is higher on autosomes relative to the X chromosome, this may be best explained by ancestral population declines (Van Belleghem *et al*., 2018). Given these predictions, our results are consistent with ancestral population contractions generating confounding signals of lower levels of genetic divergence on the X chromosome compared to the autosomes in closely related species in *Goodeinae*.

The evolution of viviparity could drive diversification rates in Goodeinae (Helmstetter *et al*., 2016), and genes involved in placental viviparity could be expected to show rapid evolution (Crespi and Semeniuk, 2004). At a broad level, we find limited support for this idea. Gene ontology for all genes under positive selection in at least one internal branch in *Goodeinae* show enrichment for reproduction and cilium movement. The lack of a clear link indicated by gene ontology may be due to difficulty in detecting positive selection at short branches where incomplete lineage sorting is pervasive. We also surveyed genes under positive selection across multiple branches and found a number of canonical immune response genes including interleukin-2 receptor subunit gamma (*IL-2RG)*, an essential constituent signalling component of many interleukin receptors (Russell *et al*., 1993); cluster of differentiation 4 (*CD4*), a central component of adaptive immune response (Luckheeram *et al*., 2012); and Lymphocyte activation gene 3 (*LAG3)*,a gene important in modulating adaptive immune responses (Grosso *et al*., 2007). Positive selection on immune response genes has been shown to be a relatively common feature of surveys of positive selection across mammals, birds, and invertebrates (Jiggins and Kim, 2007; Kosiol *et al*., 2008; Shultz and Sackton, 2019). We also find evidence of introgression of genes involved in adaptive immune response between *A. toweri* and an ancestral lineage leading to the divergence of *Ameca*, *Xenoophorus, Goodea, Xenotaenia* and *Ilyodon*, which, taken with evidence of positive selection, may indicate adaptive introgression of genes involved in MHC II pathway. Recurrent positive selection on immune-related genes, particularly genes involved in MHC II pathway, have also been associated with the evolution of viviparity in pipefish and seahorses (Roth *et al*., 2020), and broadly across Cyprinodontiformes (Yusuf *et al*., 2022).

In specific branches within *Goodeinae*, we identified genes known to be important in viviparity that were under positive selection. For example, we find positive selection in *ar* and *fshr* well known mediators of hormone synthesis and regulation in mammalian viviparity and reproduction. We also find evidence of positive selection on *igfr1*, but not *igfr2*, where parent-of-origin expression has been shown to mediate drastic changes to offspring size and positive selection has been observed in placental fish (DeChiara, Efstratiadis and Robertsen, 1990; DeChiara, Robertson and Efstratiadis, 1991; O’Neill *et al*., 2007). Saldivar Lemus *et al*. (2017) show similar parent-of-origin methylation effects on *igf2* are also present in *G. multiradiatus*, suggesting that insulin-like growth factor genes mediating sexual conflict over offspring size may be under rapid evolution within *Goodeinae*.

Additionally, our phylogenetic analysis suggests an evolutionary reversal in the form of trophotaenia reversal occurred in *A. toweri*. In trying to understand the genes that may have been involved in this, we inferred positive selection in the terminal branch leading to *A. toweri* and also identified genes that were expressed in the trophotaenia of *G. multiradiatus* (Du *et al*., 2022) and looked for loss of function mutations in these. Whilst we found no significant enrichment of any gene ontology terms, we found some genes involved in fin and appendage development, suggesting a potential link between genes identified using our approach and trophotaenia reversal in *A. toweri*. However, genes implicated in trophotaenia reversal may also be under relaxed selection leading to elevated gene-wide divergence (Hiller *et al*., 2012; Wertheim *et al*., 2015).

## Conclusion

Geological and climactic changes can have profound effects on demography, often resulting in complex speciation histories. The *Goodeinae* radiated rapidly during the Miocene and experienced constant fluctuations in population size throughout divergence, owing to periodic volcanism. We found evidence that ancient and contemporary expansions and crashes in effective population size have affected genetic variation genome-wide in closely related pairs. Volcanic activity and changes in species distribution likely facilitated ancient gene flow between ancestral lineages. We show that the same introgressed genetic variation is maintained in *A. toweri, and G. atripinnis, X. captivus* and *A. splendens*, suggesting a potential adaptive role for gene flow during the radiation of the group. Genes repeatedly under positive selection likely play a role in the evolution of matrotrophic viviparity, and we highlight specific genes that may be implicated in the reversal of trophotenium in *A. toweri*. This study highlights the importance of environmental factors, alongside internal factors such as viviparity and sexual selection, in driving species divergence, and lays the groundwork for understanding the evolution of viviparity in light of the complex demographic changes that have occurred in *Goodeinae*.

## Acknowledgements

LY was supported by a University of St Andrews studentship. MGR, CMG & YSL are supported by a Leverhulme research grant RPG-2020-181, by a Programa de Apoyo a Proyectos de Investigación e Inovación Tecnológica (PAPIIT) research grant (PAPIITIN210718) and by a Consejo Nacional de Ciencia y Tecnología (CONACyT) Ciencia de Frontera research grant A1-S-33467. PT and bioinformatics and computational biology analyses were supported by the University of St Andrews Bioinformatics Unit (AMD3BIOINF), funded by Wellcome Trust ISSF award 105621/Z/14/Z. Additional HPC (Crop Diversity) were awarded as part of a BBSRC 18ALERT grant (BB/S019669/1).

## Data availability

Scripts used to generate the assemblies and annotation can be found here: https://github.com/peterthorpe5/fish_genome_assembly. Scripts used in the comparative analysis of convergent evolution can be found here: https://github.com/LeebanY/GoodeidsPhylogenomicsComplexDemography.

Genomes and whole-genome sequencing reads are available at the NCBI under BioProject PRXXXXXXXXX.

## Supplementary Figures

**Supplementary Figure 1:**
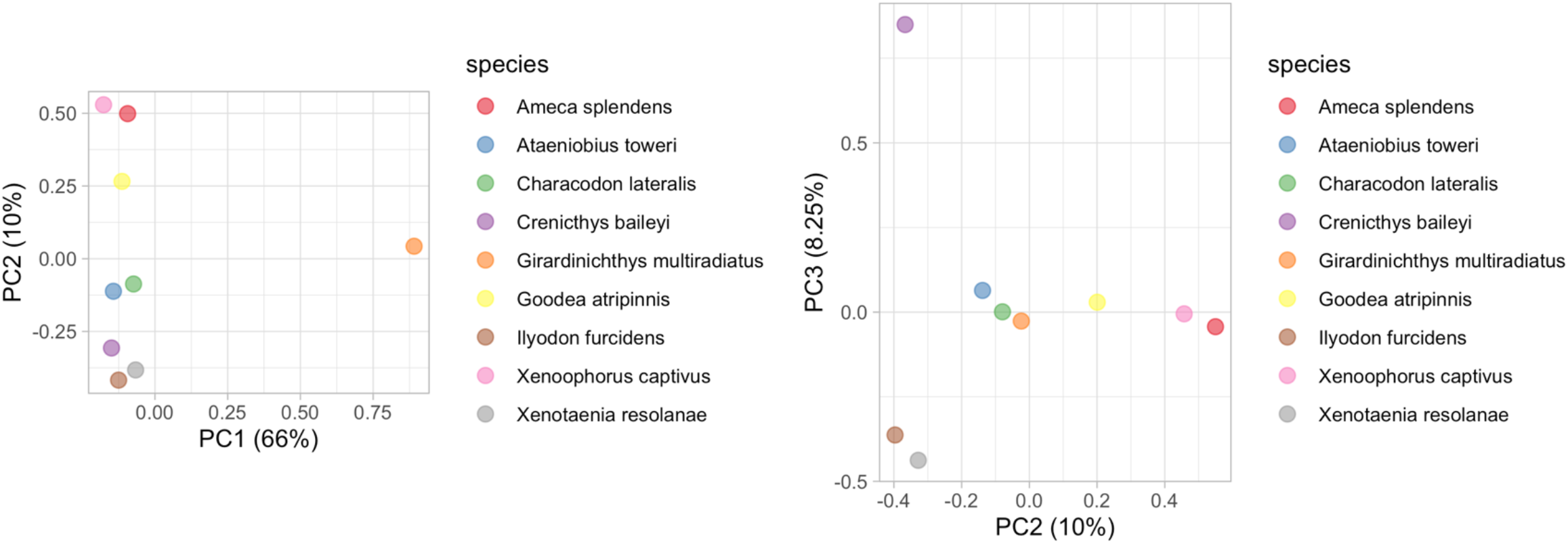
Principal component analysis (PCA) of pruned whole-genome data for all sampled species within *Goodeidae*. Axes represent PC axes explaining most variation in the data. Coloured data points in PCA correspond to species shown in legend.

**Supplementary Figure 2:**
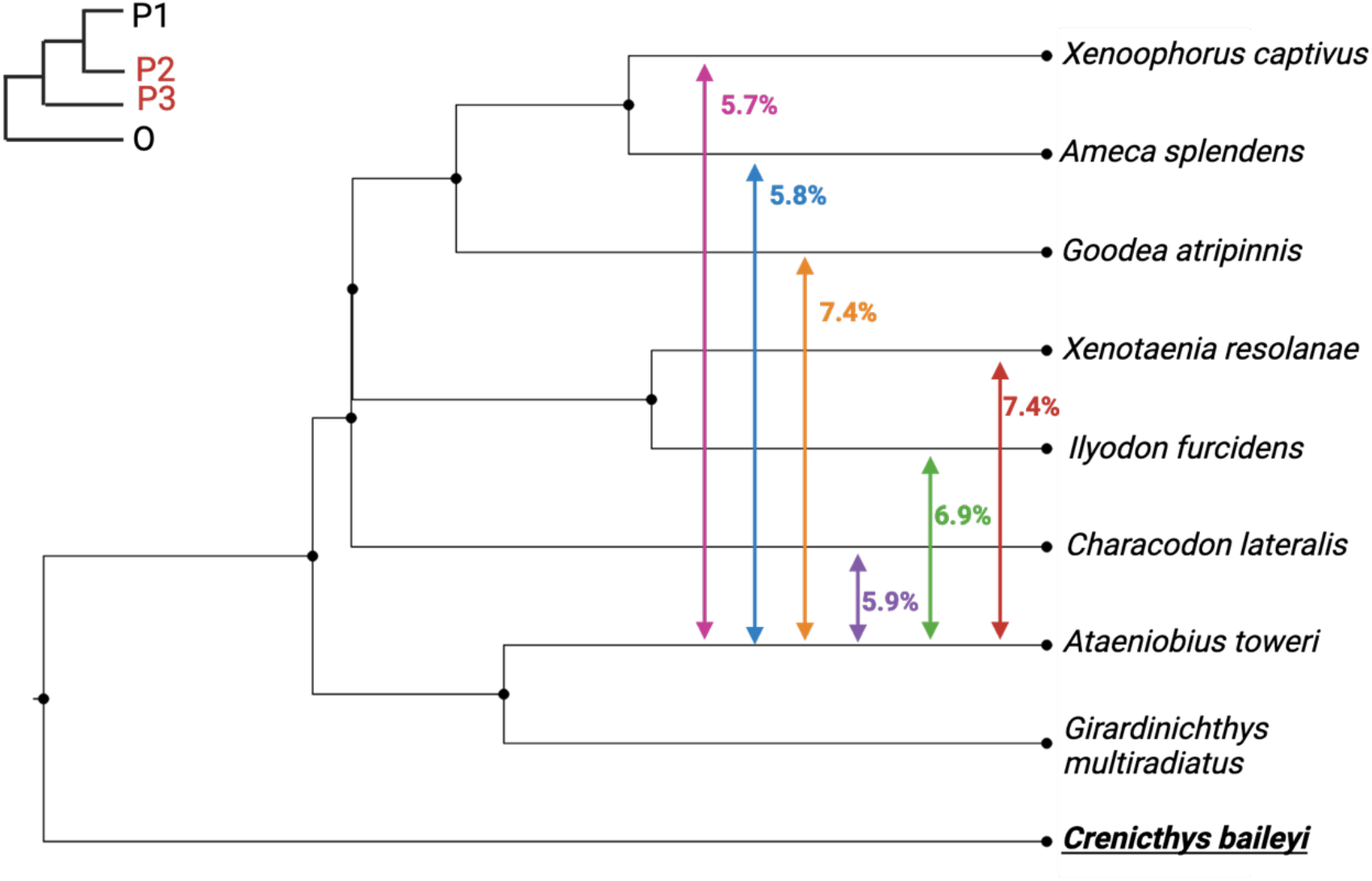
Mean genome-wide admixture proportions (f_d_) between *A. toweri (P2)* and other Goodeines (P3). *Crenicthys baileyi* used as outgroup species in all contrasts and *G. multiradiatus* was used an in-group species (P1). Phylogeny created using Figtree.

